# Determination of nuclear position by the arrangement of actin filaments using deep generative networks

**DOI:** 10.1101/2021.11.14.467997

**Authors:** Jyothsna Vasudevan, Chuanxia Zheng, James G. Wan, Tat-Jen Cham, Lim Chwee Teck, Javier G. Fernandez

**Affiliations:** Engineering and Product Development, Singapore University of Technology and Design, Singapore, 487372; Department of Biomedical Engineering, National University of Singapore, Singapore, 117576; School of Computer Science and Engineering, Nanyang Technological University, Singapore, 639798; Engineering Systems and Design, Singapore University of Technology and Design, Singapore, 487372; Mechanobiology Institute, National University of Singapore, Singapore, 117411; Institute for Health Innovation and Technology, National University of Singapore, Singapore,117599

**Author notes:** **Corresponding author** Correspondence to Javier G. Fernandez. These authors contributed equally to this work (Jyothsna Vasudevan, Chuanxia Zheng).

## Abstract

The cell nucleus is a dynamic structure that changes locales during cellular processes such as proliferation, differentiation, or migration^1^, and its mispositioning is a hallmark of several disorders^2,3^. As with most mechanobiological activities of adherent cells, the repositioning and anchoring of the nucleus are presumed to be associated with the organization of the cytoskeleton, the network of protein filaments providing structural integrity to the cells^4^. However, this correlation between cytoskeleton organization and nuclear position has not, to date, been demonstrated, as it would require the parameterization of the extraordinarily intricate cytoskeletal fiber arrangements. Here, we show that this parameterization and demonstration can be achieved outside the limits of human conceptualization, using generative network and raw microscope images, relying on machine-driven interpretation and selection of parameterizable features. The developed transformer-based architecture was able to generate high-quality, completed images of more than 8,000 cells, using only information on actin filaments, predicting the presence of a nucleus and its exact localization in more than 70 per cent of instances. Our results demonstrate one of the most basic principles of mechanobiology with a remarkable level of significance. They also highlight the role of deep learning as a powerful tool in biology beyond data augmentation and analysis, capable of interpreting—unconstrained by the principles of human reasoning—complex biological systems from qualitative data.

## Main

Quantitative research methods involve measuring some predefined features of a representative population, gathering numerical data relative to such features, and statistically analyzing the data so that it may be generalized to a larger population or explain a particular phenomenon. While the aim of statistical analysis is to remove human bias from the scientific method, feature selection is a purely human process. As researchers, we select those features that can be measured, deem important, or believe useful in hypothesizing research outcomes. There is, however, an implicit preselection, because the possible features are in all cases limited to those we can define or at least conceptualize. In other words, we cannot measure what does not exist for us.

As the selection of measurables is a rational process, the scientific method has hitherto been unavoidably constrained by human interpretation and reasoning^5^. However, this has recently begun to change. Some deep neural networks trained on large datasets are known to develop an intrinsic understanding of images in a way that goes well beyond low-level features, capturing aspects that may not be obvious or conceptualizable in our interpretation of such images.^6^ Here, we use that understanding developed by generative networks to interpret raw images of the arrangements of actin filaments in mammalian cells and demonstrate their relationship with the position of the nucleus. This correlation is commonly understood—or intuited—to exist since the mechanical interplay of both structures is known to have a major role in cell activities^7–9^ and fate^10^, and their relative misplacement is a characteristic of cell malfunction and disease^11,12^. However, demonstrating it with statistical significance is limited by the impossibility of parameterizing the spaghetti-like arrangements of cytoskeletal fibers^13^. To avoid such parameterization, we shifted the analysis from the traditional measurement of the distinct features of each substructure (i.e., actin filaments and nuclei) to the isolation of all information related to each substructure in disjoint datasets and the use of the deep generative network to find, without manually created labels or human supervision, a deterministic relationship between the different information sets.

The overall structure of the experimental design is presented in **Fig. 1**. To build the paired datasets of nuclei and cytoskeletal fibers, the two substructures were fluorescently tagged with SyTOX Deep Red (660/682) and Alexa-Fluor 488 (490/519), respectively. The selection was based on the lack of overlapping absorptions at the primary emitting wavelengths of helium–neon (HeNe; 633 nm) and argon (Ar; 488 nm) lasers, avoiding any possible crosstalk between fluorophores. Altogether, 4,900 sets of paired images at a resolution of 300 pixels per inch were taken, each set containing an average of 20 cells. The paired dataset was randomly divided into training (80%) and test (20%) images. To find the nuclear position for a given cytoskeletal arrangement, we used a transformer-based architecture^14–16^ based on the TFill network^17^. The architecture can be logically divided into three parts: (i) an encoder that takes the image of the cytoskeleton and successively embeds the two-dimensional (2D) image into high-dimensional, low-resolution feature representation; (ii) a transformer utilizes those high-dimensional features to model their dependencies with the high-dimensional features of the nuclei; and (iii) a decoder extracts those learned features in the high-dimensional space and transforms them to an image of nuclei (i.e., back to the low-dimensional, high-resolution space of common images). The results obtained were fed into a discriminator, which evaluated the proximity of the generated nuclei image to the real image and sent the results of that evaluation back to the network, training it further to improve the quality of the generated images. The auxiliary discriminator and the main generator stage a two-players-game, where two networks are trained simultaneously to compete against each other, one to generate increasingly realistic data (i.e., nuclei generation) and the other, the discriminator, improving its ability to differentiate real and generated data^18^. The iteration of this process enabled the network to identify relevant high-dimensional features in the cytoskeleton, enabling successful generation of the associated nucleus. During this process, the network was trained with qualitative data only, in the form of raw microscope images, without interpretation of the images, feature selection, or parameterization. The trained network was then used to generate the nuclei images corresponding to the test images of actin filaments. Thereafter, the images were segmented automatically, extracting the information on the number of nuclei and their diameter and position, and that information was compared with those of the real, or ground truth, nuclei.

**Fig.1.**
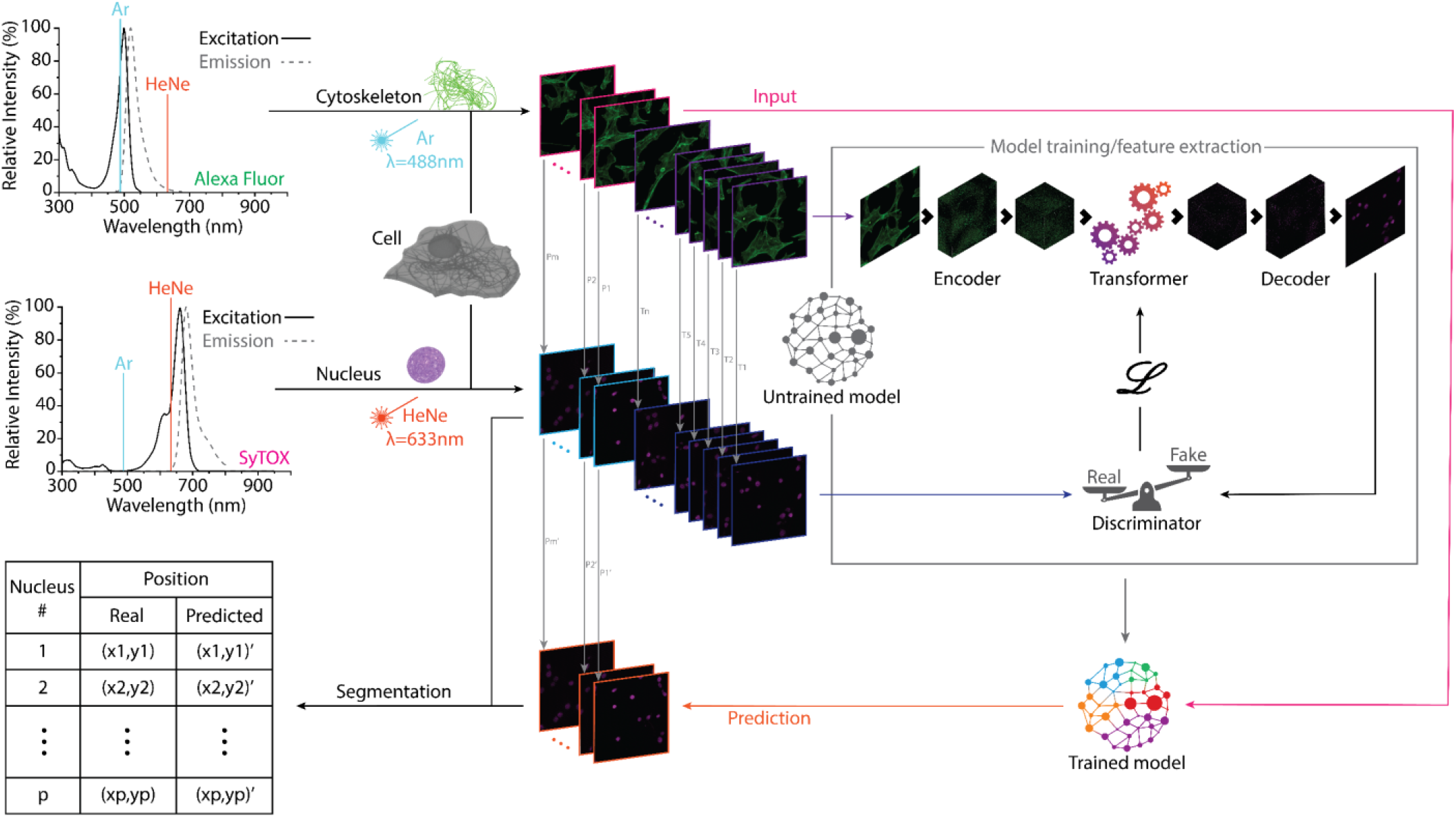
Demonstration of a correlation between arrangements of actin filaments and nuclear position. Actin filaments and nuclei information was isolated using non-overlapping fluorophores (Alexa Fluor 488 and SyTOX Deep Red). Then, 80% of the dataset of filaments was used as input to train a transformer-based network using the corresponding paired nuclei to evaluate the proximity to the real solution. The process iteration resulted in a fully trained network that was then used to generate the nuclei of the remaining 20% filament images of the dataset. The generated nuclei and their real counterparts were identified, and the coordinates of their centroids were determined to evaluate the network’s ability to predict the nuclear position using only actin filament arrangements.

The distinctiveness of the network we developed compared to other deep neural networks for image processing is that our objective was not the production of visually convincing images but of images with physiological significance, enabling the demonstration of dependencies between cellular substructures. Therefore, we eliminated image-refining steps aimed at improving appearance, common in other applications, using, for example, only a TFill-Coarse for image-to-image translation during training and testing. The encoder included a block of residual networks (ResNet, **Fig. 2a**), enabling a fast and smooth flow of information across the network by avoiding training for irrelevant layers (i.e., not adding accuracy to the outcome)^19^. The encoded vectors obtained from the input images were then fed into the transformer layer **(Fig. 2b)**, whose focus was on accessing the long-range information related to the fibrillar organization in the entire cell. Using a self-attention mechanism, the transformer ensured that all regions of the image, regardless of their location relative to a nucleus, had equal opportunities of flowing through the network’s layers^17,20^. In this way, we prevented the network’s bias toward finding an agreement between the predicted information and its immediate surrounding (preferred when filling in missing information in photorealistic images)^21^ and encouraged agreement of the predicted nuclei with all information related to the fibrillar configuration. Finally, the feature maps were projected back into completed high-resolution images by the decoder and its upsampling layers.

**Fig.2.**
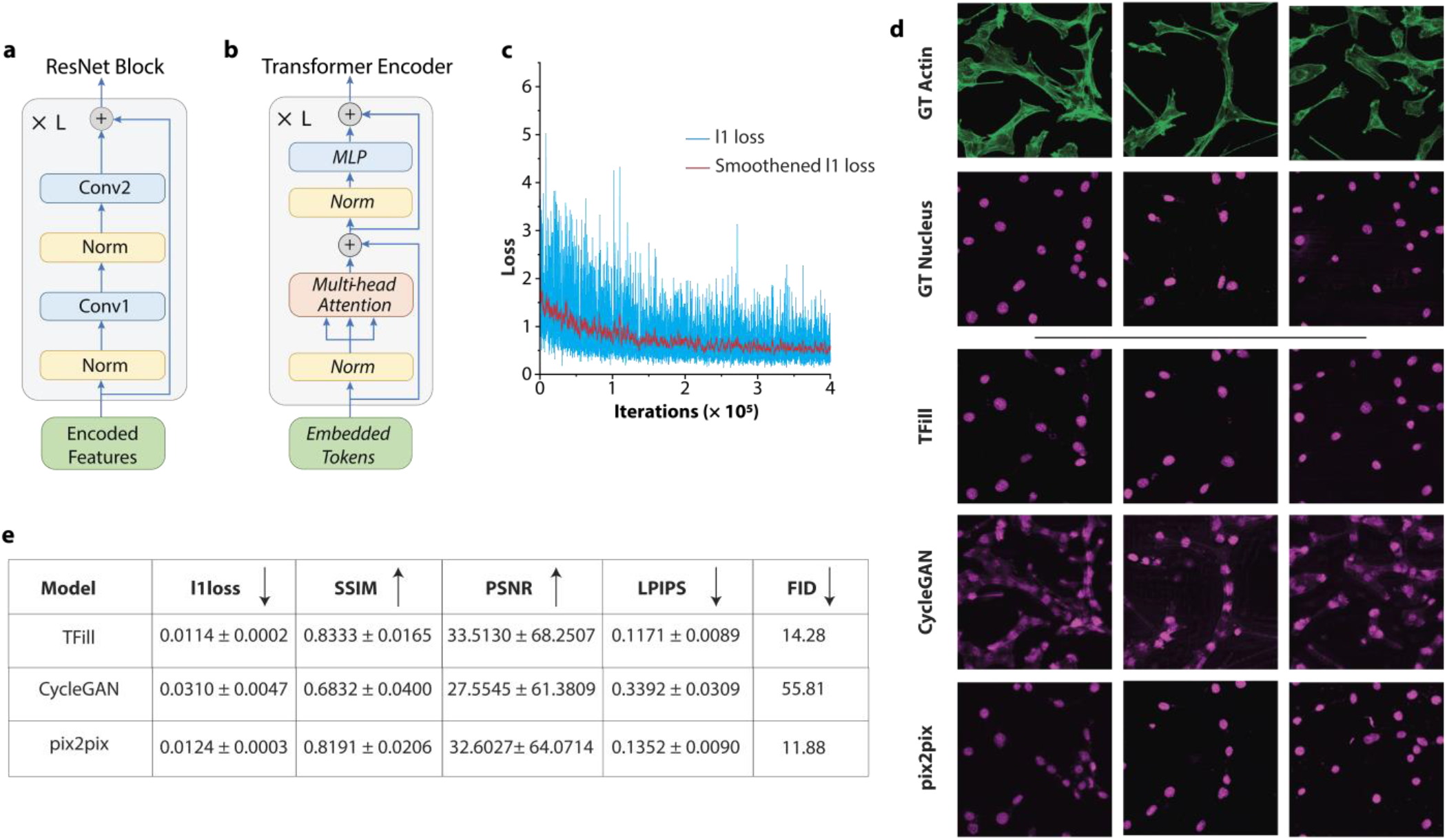
Assessing the performance of TFill network and comparison with state-of-the-art generative models. **a,** Encoder and Decoder layers comprise of a traditional convolutional neural network (CNN)-based ResNet block. **b,** The detailed architecture of the transformer encoder with self-attention mechanism. **c,** Reconstruction loss convergence as a function of iterations. **d,** Visual comparison of TFill generated nuclei images with trending image translation models. **e,** Quantitative comparison of TFill generated images with other image translation models using various metrics from computer vision (↓ Lower is better; ↑ Higher is better)

We used two distinctive loss functions during the training process. Since the targeted results, namely the nuclei, were relatively small compared to the background, we used a weighted reconstruction loss to correct the imbalance between the extensive dark background and the small, fluorescent nuclei.^22^ This decision was taken after our initial attempts to predict nuclei, which were strongly biased toward the generation of black images, deemed “realistic” by the discriminator because of their proximity to the (mostly black) ground truth. By increasing the weight of the nuclei pixels (by an order of magnitude) in the calculation of the reconstruction loss, the system was encouraged to produce nuclei to achieve acceptable levels of “realism.” Then, an adversarial loss function^18^ was used to evaluate the proximity of the generated nuclei images to real images of nuclei. This loss function was used and continuously refined by a discriminator trained to spot “fake” (i.e., generated) images of nuclei by comparison with the real images of nuclei. Concurrently, the generator was attempting to fool the discriminator by generating more realistic images while simultaneously minimizing the improving loss function.

The network training converged after 10^5^ iterations **(Fig. 2c)**. The trained network was then used to generate the nuclei of the testing dataset. The photographic characteristics of the results were evaluated on multiple metrics, consisting of pixel-level ℓ1 loss curves, region-level Structural Similarity Index (SSIM)^23^, and Peak Signal-to-Noise Ratio (PSNR)^24^, image-feature-level Learned Perceptual Image Patch Similarity (LPIPS)^25^, and dataset-feature-level Fréchet Inception Distance (FID)^26^ (**Supplementary Methods 1.7**). Despite the network architecture, which was designed for biological significance to the detriment of photorealism, the generation of images by the developed TFill network was on a par with those generated by state-of-the-art image translation networks geared toward realism, such as CycleGAN and pix2pix (**Fig. 2d and e**)^27,28^. It was not surprising with the structured paired dataset we used that the TFill-based network outperformed CycleGAN, which is conceived for mapping unpaired datasets. More interesting was the comparison with pix2pix, which is also explicitly designed to map paired images but focuses on photorealism^27^. Despite the different aims, both networks produced comparable metrics and realistic images. However, the TFill-based network showed a consistent output irrespective of the number of nuclei in the image, while pix2pix often generated noisy images and a recurrent “mosaic” artifact. Both effects were more frequent in images with many nuclei and were very likely introduced during the upscaling of the images to improve their quality. The performance of TFill **(Supplementary Figure 1)** in generating photorealistic images, outdoing networks specifically built for this purpose, highlights the possible use of the presented architecture beyond the aim of this study, namely for predicting fluorescent labels, significantly reducing the processing time of biological samples^29^.

In a further step, we evaluated the ability of the developed network to produce scientifically significant data, rather than just realistic images. We extracted the nuclear information using an automated segmentation system to determine and characterize the nuclei of the generated and real nuclei images. We also manually characterized a subset of 120 paired images as quality control for the automated characterization of the complete dataset. Overall, similar results were observed when the counting was done by automatic segmentation and matching or by manual analysis (**Fig. 3a**). In the dataset of real images, 9,659 nuclei were isolated and characterized automatically, while 8,151 (84%) were detected on the synthetic dataset. Similarly, we manually counted 1,359 and 1,069 (i.e., 79%) nuclei in the ground truth and generated datasets, respectively. The 5% difference between automated and manual counting resulted from the different thresholds of complete nucleus used by a human and by the algorithm when counting nuclei at the edges of the images; whereas the human took an educated decision on when to consider a nucleus to fall within an image, the segmentation algorithm tended to detect and count all nuclei partially falling outside the image, resulting in additional counts. On the other hand, the approximately 20% difference between the total real nuclei and those generated by the network resulted from the independent sources of information used to produce the images of ground truth nuclei (i.e., from staining) and generated nuclei (i.e., from actin fibers). Those cases where the information on the actin fiber database was poor or missing resulted in a missing generated nucleus, while the real counterpart was properly stained and identified. This resistance of the network to produce nuclei without enough fiber information is a consequence of the deliberately conservative design of the network, which prioritized quality over quantity, in contrast with the usual “aggressive” approach of image-to-image translation neural networks, where finding a solution (or several) is the priority.

**Fig.3.**
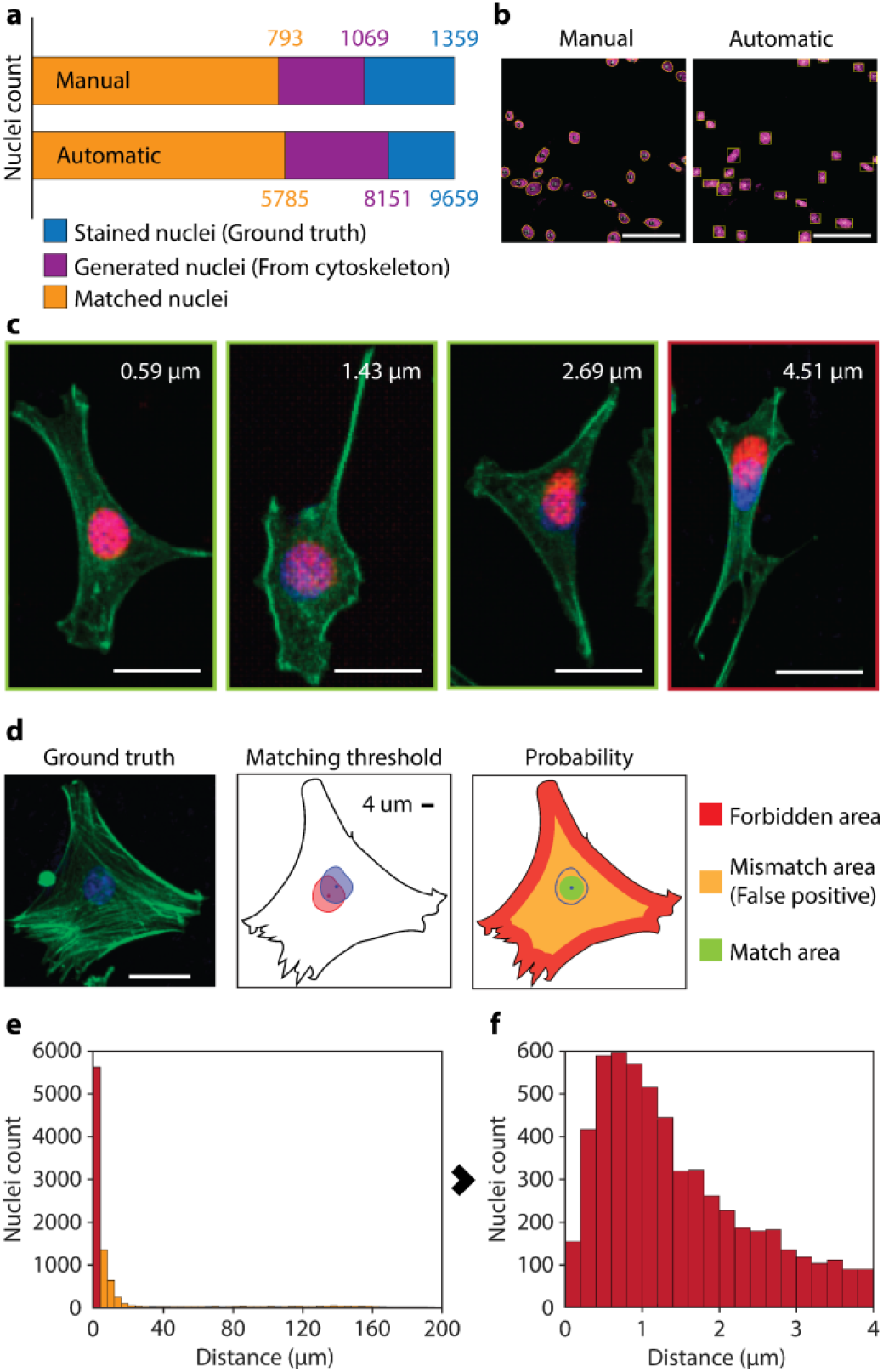
Positioning of nuclei generated from arrangements of actin filaments. **a,** Results of the identification and matching of real and generated nuclei by an automatic counting of the whole generated dataset and by manual counting of a subset of images. Stained nuclei refer to those recorded directly using fluorescent microscopy. Generated nuclei are those produced by the neural network using actin filament arrangements. Matched nuclei are those generated at less than 4 μm of its real counterpart. **b,** Manual (left) and automatic (right) processing of the same image. In manual processing the profile of the nuclei is drawn to calculate the centroid and the nuclei matched by comparison with the real counterpart. On the other hand, the automatic processing automatically identified the nuclei and generated their bounding boxes, matching generated and real nuclei based on the maximization of the overlapping areas of the bounding boxes. **c,** Several examples of generated nuclei (red) and their corresponding real nuclei (blue). The first three images (green frame) correspond to nuclei generated within the average nuclear radius (4 μm) from their real position. The last image (red frame) corresponds to a mismatch, where the generated nucleus is too far from its real position (See Supplementary Fig.2 for full images; Bars are 20 μm). **d,** Example of a cell and the relative distance of 4 μm within the cytoplasm. The probability of randomly positioning the nucleus within the cytoplasm can be identified as the ratio of possible matched positions for the centroid (green area) with respect to all possible positions (orange area). Those possible positions of the centroid located at less than the nuclear radius from the edges of the cell (red area) are discarded under the premise that the nucleus cannot be positioned partially outside the cell (See Supplementary Fig. 3 for further analysis of probabilities; Bar is 20 μm). **e,** Distribution of the distances of the generated nuclei respect their real position. 71% of the nuclei are situated at less than 4 μm of their real position. **f,** Distribution of distances of the generated nuclei considered matched (<4 μm). 40% of the matched nuclei are located at less than 1μm from their real position.

We then demonstrated the deterministic relationship between the nuclear position and the arrangement of actin filaments. We matched each generated nucleus with its real counterpart and calculated the Euclidean distance between their centroids. This was performed automatically by producing a bounding box for each identified nucleus, calculating the overlapping ratio (OR) of each bounding box of a generated nucleus with all those of real nuclei, and pairing them by maximizing the OR (**Fig. 3b**). As before, we kept the manually processed subset as control. The bounding boxes were used to also calculate the centroid of each nucleus, which, in the case of generated nuclei, depend on the ability of the neural network to predict the right position and, to a lesser extent, the shape of the nucleus (**Fig. 3c, Supplementary Fig. 2**)^30^. The neural network, using only cytoskeletal information, positioned 71 ± 1% (for a confidence interval (CI) of 95%) of the generated nuclei within the radius of the real nucleus (4 μm), and almost one out of three nuclei (29 ± 1%, for a CI of 95%) were generated at less than 1 μm from the center of the real nucleus (**Fig. 3d and 3e**).

The consensus for biological experiments is to discard the null hypothesis for a p-value of < 0.05. In our experiments, the extreme level of confidence makes p values meaningless^31^ (**Supplementary Methods 1.9)**. For example, given that the system has one out of 500 chances to randomly place the nucleus’s centroid at less than 4 μm of the correct position within the image (159.41×159.41 μm), the correct localization of 71% of the nuclei corresponds to a p-value of approximately 10^−2170^. Focusing the analysis on the location of the nucleus in the cell, rather than in the image, we would be assuming that the neural network somehow achieved the correct localization of the nuclei by developing: i) an understanding of the low-level characteristics of the filament arrangements (a feature conceptually similar to the limits or shape of the cell); ii) the understanding that the nuclei must be within those limits; and iii) the skills to predict the size and shape of the nuclei (**Fig. 3d**). In such a situation, the problem would reduce in its last instance to the successful localization of the nucleus within the cytoplasm, a task that, in the most optimistic situation, can be randomly achieved in one out of two cases (**Supplementary Fig. 3**). This restrictive situation results in an equally negligible p = 6.6×10^−130^, yielding strong evidence against the null hypothesis (i.e., a random positioning of the nuclei within the cytoskeleton) and demonstrating, with overwhelming significance, one of the most basic principles of cell biology: the correlation between the position of the nucleus and the actin filaments.

In sum, to demonstrate the correlation between the position of the nucleus and the cytoskeleton in cells, we isolated the information about the substructures using non-overlapping fluorophores and laser lines. We developed an image-to-image translation algorithm based on a TFill network and trained it, using unparameterized images of actin filaments, to extract high-dimensional features relatable to nuclear information. The network’s success in accomplishing its task was measured by predicting several thousand nuclei from the arrangements of actin filaments. Seventy-one per cent of the nuclei were generated within the surface of the real nuclei, and almost half of those at less than 1 μm distant from their real position, demonstrating with astounding significance the hypothesis of a deterministic relation between the arrangements of the actin filaments and the position of the nucleus. This demonstration illustrates the ability to use deep neural networks, outside data analysis or augmentation, as a method to interpret reality beyond the limitations of human conceptualization, and, specifically, to extract features of systems with a complexity unsuitable for quantitative parameterization. Our results also evidence the conditions enabling a transition from a methodology in biology based on human analysis to methods of data acquisition focused on curating information for non-human interpretation.

## Supplementary Information

The Supplementary Information contains a detailed description of the materials and methods used and their discussion, three additional figures on the performance of the generative network, the generation of nuclei, and the statistical analysis.

## Acknowledgments

The Singaporean Ministry of Education has supported this research through the MOE2018-T2-2-176 grant to JGF.

## Contributions

JV and JGF conceived the idea of using machine learning to demonstrate the correlation between actin filament arrangement and nuclear position and designed the experiment. JV, CZ, and JGF conceived the idea of using a deep image-to-image translation network for the task. CZ developed the network and performed the training with the help of JV and the supervision of TC. JV developed and performed the automated and manual segmentations to extract nuclear information. JV prepared and collected the biological data with the supervision and advice of LCT. JGW, JV, and JGF worked out the statistics. JV, CZ, and JGF wrote the manuscript with inputs from all the other authors.

