## Supplementary Materials and Methods, and Figures for "Determination of nuclear position by the arrangement of actin filaments using deep generative networks"

This material includes:

**1 Methods**

**1.1** Cell culture
**1.2** Fluorescence labeling
**1.3** Imaging
**1.4** Dataset preparation
**1.5** Neural network – architecture
**1.6** Loss functions
**1.7** Neural network – Evaluation
**1.8** Comparison of nuclei count and positioning in real and generated images
**1.9** Distribution and probability of the nuclei position within the image and the cell

**2 Figures**

**Figure 1** TFill Network performance assessment
**Figure 2** Generation of nuclei and matching
**Figure 2** Nucleus distribution in the image and cell

**3 References**

### 1 Methods

#### 1.1 Cell culture

Mouse fibroblasts (NIH/3T3) (ATCC) were cultured and maintained in Dulbecco's Modified Eagles Medium (high glucose DMEM, Nacalai Tesque) supplemented with 10% (v/v) Fetal Bovine Serum (FBS, Life Technologies) and 1% (v/v) Penicillin-Streptomycin mixed solution (Nacalai Tesque). Cells were incubated in normal physiological conditions (37°C, 5% $CO_2$), passaged every three days, and their media was replenished every two days.

#### 1.2 Fluorescence labelling

Upon confluence, the cells were trypsinized, centrifuged, and resuspended in fresh DMEM media at a concentration of $1 \times 10^6$ cells/ml. The cells were seeded on cover glass substrates (22 × 22 mm, thickness = 100 µm) at a seeding density of 20000 cells per substrate to ensure adequate spacing between the cells. All substrates were maintained in standard cell culture conditions (37°C, 5% $CO_2$) for 24 hrs. The cell-seeded glass substrates were rinsed with Dulbecco's Phosphate Buffered Saline (D-PBS) and fixed with 4% (w/v) Paraformaldehyde (PFA) (Merck Millipore) solution in D-PBS for 15 min. Samples were then washed three times (3X), 5 min each time, with D-PBS. Cells were permeabilized with 0.5% (v/v) Triton X-100 in D-PBS for 10 min and then washed 3X, 5 min each time, using D-PBS. Subsequently, the cells were blocked with 3% (w/v) bovine serum albumin (BSA, Gold Biotechnology) for 1 hr. Samples were stained with a 1:500 (v/v) dilution of Alexa-Fluor® 488 phalloidin (Life Technologies) for 30 min and washed 3X, 5 min each time, with D-PBS. Nuclei were stained with a 0.1 µM SyTOX™ Deep Red Nucleic Acid stain (Life Technologies) for 10 min, following which, the substrates were washed thoroughly with D-PBS to remove unbound stains. The coverslips were mounted (Fluoroshield mounting media, Abcam) and stored in the dark until further use.

#### 1.3 Imaging

The images were acquired using a confocal microscope (Zeiss LSM 710) equipped with a 40× lens with numerical aperture 0.95 connected to Zen Black. Coverslips containing fixed cultured cells are mounted on top of a motorized stage. The microscope was programmed to acquire Z-stack images from a 25 x 25 square tile region (5.250 x 5.250 mm), thus acquiring a total of 625 image fields. The motorized stage can translate across x and y directions. The total thickness of each Z-stack was set to 20 µm, and images were acquired at an interval of 1 µm for each field. Two independent channels were acquired: Alexa Fluor 488 Phalloidin (Excitation (max): 495 nm, Emission (max): 518 nm) and SyTOX Deep Red (Excitation (max): 660 nm, Emission(max): 682 nm). During image acquisition, it was ensured that a stitching algorithm was not employed to facilitate the splitting of the large image area. A total of 4900 image pairs were collected across eight samples.

#### 1.4 Dataset preparation

All acquired images were processed using Zen Blue Lite software. Images were first subjected to maximum intensity projection (MIP) to convert the 3D data into a single 2D image. It was ensured that all the features were visible in the single image plane. The large image area was split into two separate channels (Actin and Nuclei) and small, square image tiles, each of size 210 x 210 µm. The images were arranged into folders, each containing one Actin-Nucleus pair image data, using a custom-written Python script. 4900 image pairs were obtained and randomly divided into training (80% of the total dataset) and testing (20% of the total dataset) images.

#### 1.5 Neural network – architecture

Our network architecture extends the TFill network from Zheng *et al*. We only used TFill-Coarse for image-to-image translation since this study focuses more on nuclei positioning than the realistic appearance generation[1]. Its architecture can be logically grouped into three parts: (i) **an encoder** that takes an image $I \in \mathbb{R}^3$ as input and then successively embeds the 2D image into high-dimensional latent space thus yielding a low-resolution token representation $z_0$; (ii) **a transformer** that captures long-range dependencies between encoded token representation and then outputs the global feature representation; (iii) and **a decoder** that takes the learned feature representation and generates all nuclei based on cell shapes. It gradually upsamples the low-resolution feature maps to high-resolution feature maps to achieve the original resolution images.

The transformer architecture was firstly introduced in Natural Language Processing (NLP) and later widely used in various computer vision (CV) tasks, such as scene classification, object detection, instance segmentation, image generation, and translation. A transformer encoder layer consists of a Multihead Self-Attention (MSA) and a Multilayer Perception (MLP) block. The MSA is applied to capture the long-range relationship between each token, while the MLP is responsible for further transforming the merged features from the MSA layers. Furthermore, to achieve the more complex features, the Layernorm (LN) is used before the MSA and MLP block for none-linear projection. They are expressed by:

$$z_0 = [x^1; x^2; \dots; x^N] + E_{pos} \quad (1)$$

$$z_l' = \mathrm{MSA}\big(\mathrm{LN}(\mathrm{z_{l-1}})\big) + z_{l-1} \quad (2)$$

$$z_l = \mathrm{MLP}\big(LN(z_l')\big) + z_l' \quad (3)$$

where $z \in \mathbb{R}^{N \times C}$ is the 1D sequence of N tokens $x$ with $C$ channels, $E_{pos} \in \mathbb{R}^{N \times C}$ is the position embedding. $z_0$ is the input sequence of the transformer while $z_l$ is the sequence output in layer $l$.

The components of the neural network are illustrated in detail in **Error! Reference source not found.**. Unlike the NLP that naturally treats each word as token[2] embedding for the 1D sequence, the visual image is in 2D without explicit word representation. Following the existing vision transformer[3-5] arch the CNN-based encoder embeds the 2D images as high-dimensional, low-resolution feature representations. The encoder is a ResNet-style convolutional neural network[6]. It is stacked with four repeated residual blocks. Each block consists of two convolutions, each followed by a leaky rectified linear unit with a leakage factor of 0.2 and a pixel-wise normalization. Besides, a skip connection is used to pass the information through a short path quickly. Each block is followed with a learned pooling operation with stride two to halve the resolution of the resulting feature map.

The transformer-based encoder consists of twelve transformer blocks. Each one can automatically capture the long-range dependencies between all locations in the feature maps. Furthermore, instead of the fixed weights in the CNN-based network, the transformer merges information using the input-dependent adaptive weightings, which are decided by the similarities between features. The decoder is an inverse operation of the encoder. It is applied to unfold the high-dimensional, low-resolution features into low-dimensional, high-resolution images. The decoder is also stacked with four repeated residual blocks as in the encoder. However, each block is followed with an upsampling layer to increase the resolution of feature maps. In parallel, the output layer is added after each block to get multilevel, multi-resolution outputs, where the output structure is inspired by StyleGAN v2[7]. The multi-resolution outputs allow predication errors to quickly backpropagate to the previous encoders that can stabilize and speed up the training. Finally, an auxiliary discriminator using adversarial learning is applied to improve the generated image quality further [8]. We directly adopt the discriminator architecture from the latest StyleGAN v2[7], which downsamples input images to 4×4 resolution, and then uses fully connected layers to judge them belong to “real(1)” or “fake(0)”. This encourages the generated results to match the distribution in the given data.

#### 1.6 Loss functions

**Weighted Reconstruction Loss:** We first use pixel-weighted reconstruction loss to enable changing the influence of imbalanced nuclei and background ratios in the target image. The loss is defined as:

$$\mathcal{L}_{rec} = \sum_{x \in \Omega} \{\alpha \cdot \left\| M(x) \odot \left(I_{out}(x) - I_{gt}(x)\right) \right\|_1 + \left\| \big(1 - M(x)\big) \odot \left(I_{out}(x) - I_{gt}(x)\right) \right\|_1 \}$$

where x is pixels in image domain Ω, $I_{out}$ and $I_{gt}$ are the generated nucleus image and the corresponding ground truth respectively, M is the binary mask map where nucleus regions are labeled as "1," and background pixels are labeled as "0". Thus, the first term of the equation is related to the nuclei regions' reconstruction, while the second term is for the background reconstruction. We use the L1 reconstruction loss for each matched pixel in the generated output and ground truth image.

In our images, most pixels belong to the black background. If we rebuild the original ground truth directly, the output will prefer to generate the black image on average. Therefore, we manually increase the weight of nuclei pixels using a factor $\alpha$ =10, as compared with the black background pixels. To do this, we will enforce the model biases to the nuclei regions, resulting in a balance training.

**Adversarial loss:** We further introduce the adversarial loss to encourage the generated nuclei's distribution to be closed to the nuclei's distribution in the ground truth. Following the previous adversarial learning, we model this *minimax game* using an adversarial loss given by:

$$\mathcal{L}_{gan} = \min_G \max_D E_{I_{gt} \sim P_{data}(I_{gt})} \left[\log D\left(I_{gt}\right)\right] + E_{I_{in} \sim P_{data}(I_{in})} \left[\log\left(1 - D\left(G(I_{in})\right)\right)\right]$$

where G is the generator and D is the discriminator, and $\mathbf{I}_{gt}$ and $\mathbf{I}_{in}$ are the data from target ground truth sets and input sets. During the training, generator(G) and discriminator(D) parameters are updated alternately. The D is trained to distinguish between generated and ground truth images by maximizing the loss function above. At the same time, the G tries to fool the discriminator to generate more realistic images by minimizing the above loss function.

#### 1.7 Neural network – Evaluation

**Fréchet inception distance (FID)**[9]: This metric calculates the mean and variance distance between the feature vectors for ground truth and generated images. A low FID score indicates better performance.

**Learned Perceptual Image Patch Similarity (LPIPS)** [10]**:** This evaluates the diversity of generated images compared to its ground truth, and it is a state-of-the-art metric that correlates to human perceptual similarity. A lower LPIPS score indicates that the generated image is more realistic and similar to the ground truth.

**Structural Similarity Index (SSIM)** [11]**:** This metric computes the perceptual distance between a translated image and its corresponding ground truth based on three indices: luminance, contrast, and structure. The higher the SSIM score, the greater the similarity of the two images.

**Peak Signal-to-Noise Ratio (PSNR)** [12]**:** PSNR calculates the differences in intensity between the ground truth and the generated image and is defined via the mean square error (MSE). A high PSNR score indicates that the intensity of both images is similar.

#### 1.8 Comparison of nuclei count and positioning in real and generated images

The number of nuclei and their positioning were quantified using built-in functions in MATLAB (MathWorks, Natick, MA)[13]. The images were initially subjected to binarization via HSV thresholding. The properties of the nuclei were recorded from the thresholded images, and the nuclei were counted from the **regionprops** function. The function also provides information such as area, centroid, and bounding boxes around the nuclei region. We then filtered the segmented components with a minimum area of 50 pixels to eliminate noise due to thresholding deficiencies. In addition, poorly thresholded images were manually eliminated from further analysis since the goal is not to test for segmentation accuracy. 729 images out of 980 test images were used to calculate errors in nuclei count and positioning accuracy. The percentage error in nuclei detection was computed as follows:

$$\text{Error (\%)} = \frac{|\text{Nuclei count}_{\text{GT}} - \text{Nuclei count}_{\text{Gen}}|}{\text{Nuclei count}_{\text{GT}}} \times 100$$

where Nuclei $count_{GT}$ indicates the number of nuclei in the ground truth and Nuclei $count_{Gen}$ indicates the number of nuclei in the generated image. Subsequently, for every generated nucleus, the bounding box resulting from **regionprops** function was compared with that of ground truth to compute the overlap ratio using the MATLAB in-built function **bboxOverlapRatio**. The overlap ratio (OR) is an intersection over union metric (IoU), which was computed as follows:

$$Overlap\ Ratio = \frac{Area\ (BB_{GT})\ \cap\ Area\ (BB_{Gen})}{Area\ (BB_{GT})\ \cup\ Area\ (BB_{Gen})}$$

where $BB_{GT}$ and $BB_{Gen}$ indicates the bounding boxes of ground truth and generated nuclei respectively. This is used to match generated nuclei centroids to ground truth nuclei centroids. Then, an error metric is computed as a Euclidean Distance (ED) between the two coordinates as follows:

$$\text{ED} = \sqrt{(X_{GT} - X_{Gen})^2 + (Y_{GT} - Y_{Gen})^2}$$

We also use the Clopper and Pearson approach to establish 95% confidence intervals for all of the performance metrics in this study[14].

#### 1.9 Distribution and probability of the nuclei position within the image and the cell

Statistical analysis is done by modeling the number of correct predictions made by the neural network using a binomial distribution, with a probability of success p for each prediction. Due to the very high number (n= 8151) of trials, a normal distribution is used to closely approximate the binomial. Based on 5785 correct predictions at the 4um distance, the 95% score confidence interval for p is computed to be 71.0+/-1.0%. Similarly, based on 2328 correct predictions at the 1um distance, the 95% score confidence interval for p is computed to be 28.6+/-1.0%.

To calculate the network's success in predicting the correct position of the nuclei, we first calculated how successful any system would be doing it randomly. To do so, we define the ratio of "right positions" and "possible positions" on the area where the nucleus is generated (Supplementary Fig. 3). The correct positions are those within the nucleus area, taking 4 µm as the average radius for the cells employed in this study. This assumption is made based on the possible divergence of the centroid due to imaging and the different shapes of the real and generated nuclei. We calculated the ratio within the whole image as well as within the cell. To provide a realistic approximation and avoid an overrepresentation of possible positions, we impose the additional restriction that the whole nucleus must be inside the cell for the calculation. Therefore, those possible positions situated at a distance of less than a nuclear radius from the edges of the cell were discarded. To get realistic values, we first modeled cells as a circular entity based on the average radius in a confluent culture. Then, we repeated the same calculation for the images used in this study. In that case, the position ratios are strongly dependent on the cell geometry.

We calculated the ratios and associated p-values for several cells, including the two most extreme examples found. In one end are cells narrowly spread around the nucleus and with the cytoplasm distributed in several protrusions (Supplementary figure 3C1), strongly limiting the possible positions of the nucleus to the small central space. On the other extreme are cells uniformly spread and with short or inexistent protrusions (Supplementary figure 3E), where the possible positions for the nucleus roughly cover the whole cytoplasm. We also repeated the same calculations using a threshold distance of 1 µm between the centroids of the real and predicted nuclei. In that case, the number of matched nuclei drops to almost half; however, the probability of correctly positioning it by chance also decreases dramatically, resulting in similarly low p-values.

All the p values reported in this study are negligible, and it makes no difference to have a $p=10^{-100}$ or $10^{-1000}$, since both are in a range of overwhelming statistical significance. In that situation, we find the confidence interval (reported in the main text) a much informative way to report an error. Despite the absurdity of these negligible p values, we have included them as a comparison with the current standards in biology to highlight the power of the approach presented here. P values are a standard in biological and biomedical research, fields that struggle to achieve p values $< 0.05$, a thresholds that hardly ensure reproducibility of the results [15]. Here, however, analyzing a similar system, achieve p values that in the worst scenario are $< 10^{-100}$., giving a clear picture of the quality of the demonstration compared to those currently accepted in biomedical studies.

#### 2 Figures

**a**

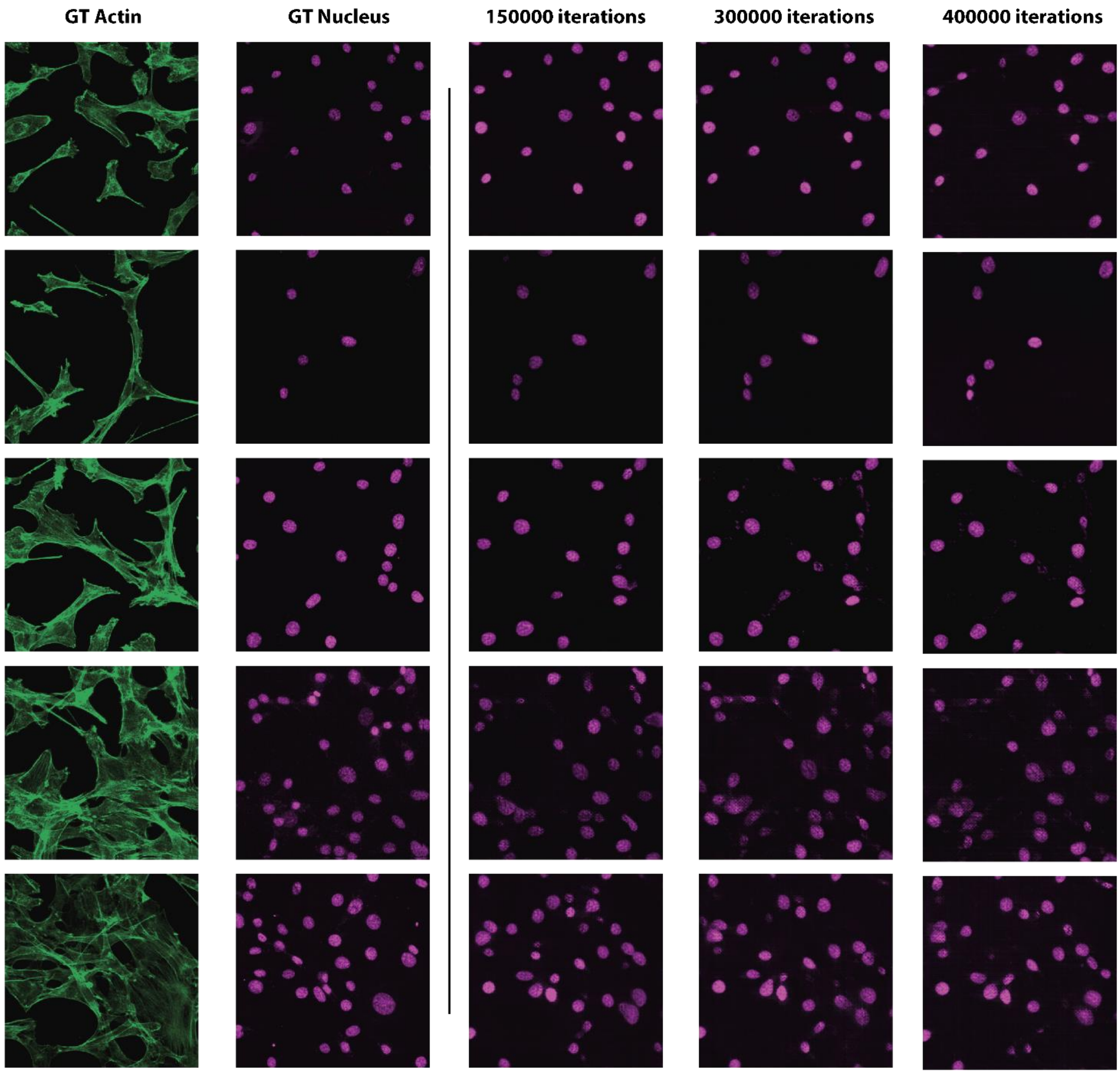


**b**

| Model | Iterations | l1loss ↓ | SSIM ↑ | PSNR ↑ | LPIPS ↓ | FID ↓ |
|---|---|---|---|---|---|---|
| TFill | 150000 | 0.0117 ± 0.0002 | 0.8333 ± 0.0175 | 32.9127 ± 65.3693 | 0.1204 ± 0.0105 | 32.77 |
| | 300000 | 0.0117 ± 0.0002 | 0.8328 ± 0.0174 | 33.1595 ± 65.2648 | 0.1209 ± 0.0092 | 21.66 |
| | **400000** | **0.0114 ± 0.0002** | **0.8335 ± 0.0165** | **33.5130 ± 68.2507** | **0.1171 ± 0.0089** | **14.28** |

**Fig. 1 | Iterative Evaluation of TFill Neural Network Performance. a,** Visual comparison of TFill generated nuclei images with altering training iterations; **b,** Quantitative comparison of TFill generated images with altering training iterations using various metrics from computer vision (↓ Lower is better; ↑ Higher is better)

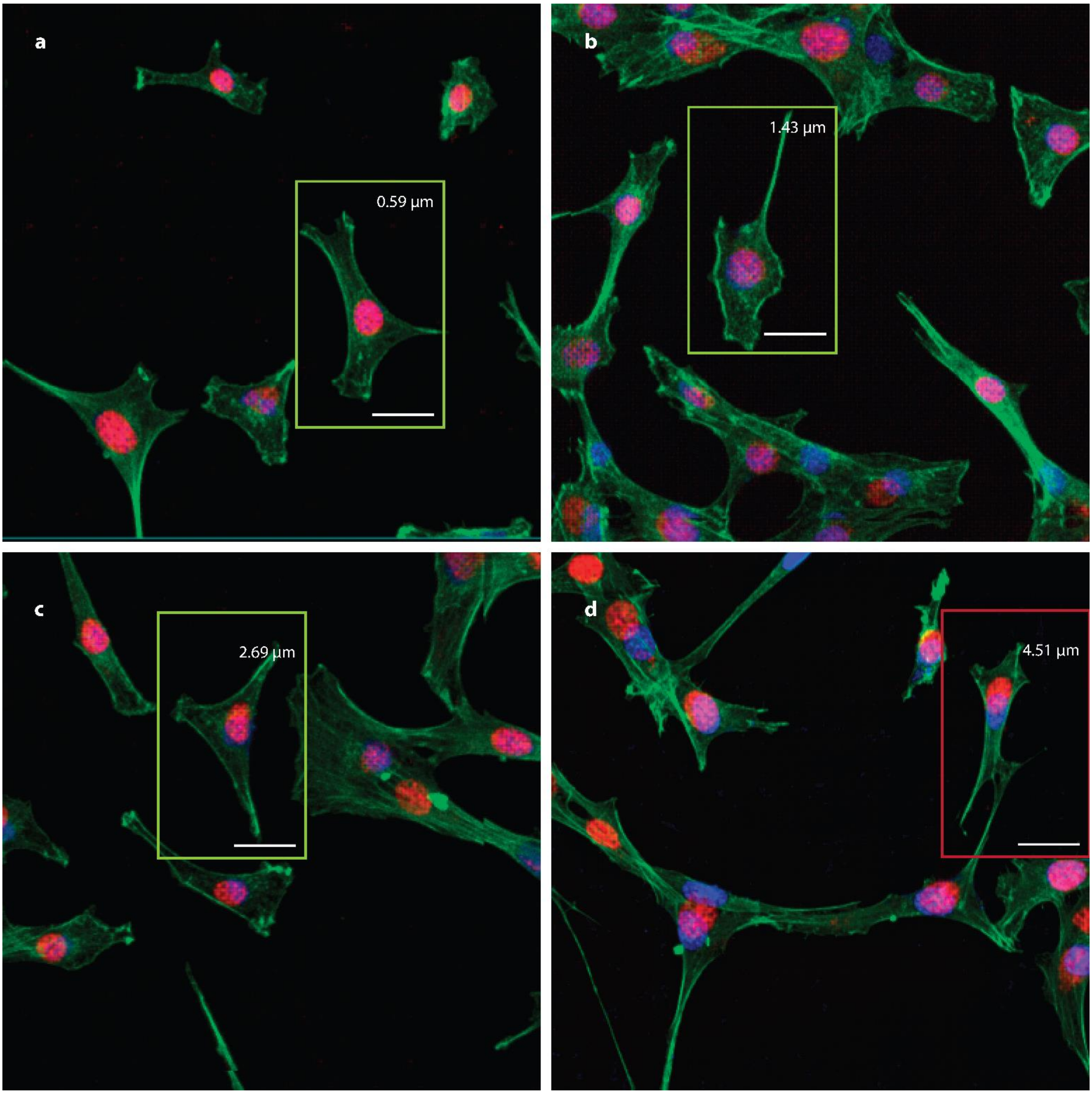


**Fig. 2| Generation of nuclei and matching.** Full images of panels in Figure 3c of the main text. The generated nuclei (red) and real nuclei (blue) have been placed over the images of the actin fibers used for their generation. Bars are 20 µm, and the values correspond to the distance between the centroids of the generated and real nuclei (i.e., ground truth) of the framed cell.

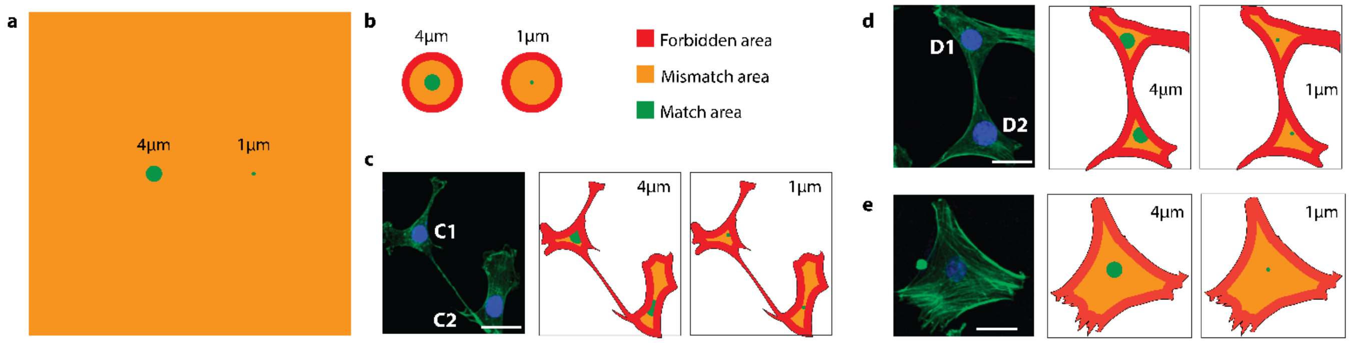


| | 4 µm (5785/8151) | | | | 1 µm (2328/8151) | | | |
|---|---|---|---|---|---|---|---|---|
| | Match | Mismatch | Z-score | p-value | Match | Mismatch | Z-score | p-value |
| A- Full image (159.41x159.41µm) | 0.001978 | 0.998022 | 1438.124 | 0 | 0.000124 | 0.999876 | 2318.232 | 0 |
| B- Modeled circular cell (Nucelus diameter 8µm, cell diameter 30µm) | 0.132231 | 0.867769 | 153.9169 | 0 | 0.008264 | 0.991736 | 276.5795 | 0 |
| C1 – Extreme narrow (top) | 0.580192 | 0.419808 | 23.69674 | 0 | 0.071678 | 0.928322 | 74.87525 | 0 |
| C2- Regular cell (bottom) | 0.110951 | 0.889049 | 172.1246 | 0 | 0.019449 | 0.980551 | 174.0065 | 0 |
| D1- Regular cell (top) | 0.29502 | 0.70498 | 82.09835 | 0 | 0.021562 | 0.978438 | 164.1266 | 0 |
| D2- Regular cell (bottom) | 0.301934 | 0.698066 | 80.19432 | 0 | 0.024732 | 0.975268 | 151.6532 | 0 |
| E- Large cell | 0.059372 | 0.940628 | 248.4612 | 0 | 0.003778 | 0.996222 | 414.7504 | 0 |

**Fig. 3| Nucleus distribution in the image and cell.** In all panels, green refers to those areas where positions the nuclei centroid is considered right (Match). Orange areas are possible positions for the centroid considered a failure (Mismatch), and red areas are those positions forbidden for the nucleus centroid for being too close to the cell edge. **a,** Distribution of possible positions for the whole microscope image. The green area are those positions at either 4 or 1µm away from the centroid of the real nucleus. **b,** Represent the distribution in a circular model of a cell using the average radius of a cell in confluence. **c-e,** Examples of the distribution of positions in cells used in this study. Cells C1 and D represent the highest and lowest ratios found. The bottom table is a calculation of the p-value for all the possible distributions. A $p\text{-value}<0.05$ is commonly considered enough to discard the null hypothesis and support the alternative hypothesis. In this case, the null hypothesis is a random distribution of the nuclei with respect to the actin fibers. The alternative hypothesis is the deterministic relation of the nucleus position with the fiber arrangement. In all cases, p ranges between $10^{-100}$ and $10^{-2200}$ and have been approximated to 0.
